# Integrated Histology and Molecular Profiling of Postmortem Human Auditory and Vestibular Organs via a Poly(Methyl Methacrylate)-Based Workflow

**DOI:** 10.1101/2025.06.04.657716

**Authors:** David Bächinger, Brock Peyton, Jacqueline Neubauer, Anbuselvan Dharmarajan, MengYu Zhu, Jennifer T. O’Malley, Venus Kallupurackal, Steven Senese, Alison Brown, Sabina Wunderlin, Susanne Kreutzer, Nora M. Weiss, Heiko Richter, Adrian Dalbert, Christof Röösli, Anja Kipar, Zsuzsanna Varga, Brigitte von Rechenberg, Sami S. Amr, Andreas H. Eckhard

## Abstract

Hearing and balance disorders are the most prevalent sensory impairments, affecting hundreds of millions worldwide, yet their underlying cellular and molecular pathologies remain poorly understood. This knowledge gap stems from the inaccessibility of the ear’s sensory organs—embedded within the temporal bone (TB), the hardest bone in the body—which cannot be biopsied in living patients without causing irreversible damage. Conventional histopathology workflows rely on postmortem en bloc extraction of TBs, followed by lengthy decalcification, celloidin embedding, and manual serial sectioning of these large specimens—a process that takes one to two years, is labor- and cost-intensive, and lacks compatibility with most modern protein, DNA, and RNA assays. Here, we present a rapid, reversible polymethyl methacrylate (rPMMA) workflow that enables advanced molecular histopathology studies on formalin-fixed, calcified TBs. Our protocol uses low-temperature (−40 °C to +4 °C) resin embedding, precision near-serial sectioning (10–50 µm) via femtosecond laser microtomy or precision diamond wire sawing, and subsequent deacrylation to fully restore tissue accessibility for high-fidelity histomorphology, multiplexed immunofluorescence, whole-genome sequencing, and in situ mRNA detection (RNAscope™) assays. Compared to the gold-standard celloidin workflow, our method reduces processing time and costs by approximately 90% while integrating equivalent histomorphology with advanced molecular assays, providing a new benchmark for multidimensional studies in human hearing and balance pathologies.

## Introduction

Hearing and balance disorders are the leading global causes of sensory impairment^1–3^, yet their underlying cellular and molecular pathologies remain poorly understood—a major barrier to new therapeutic development^4^. This knowledge gap stems from the inaccessibility of the ear’s sensory organs, which lie embedded within the skull’s temporal bone (TB)—the hardest bone in the human body—and cannot be biopsied in vivo without causing irreversible damage. Consequently, human pathology studies on hearing and balance disorders rely almost exclusively on en bloc TB specimens harvested postmortem from individuals with well-documented clinical histories^5,6^.

Human TB specimens containing the ear’s sensory structures are large (approximately 38 × 38 × 50 mm^5,7^) and anatomically complex, with delicate sensory tissues suspended in air- and fluid-filled chambers encased by densely mineralized bone. Standard histology workflows—using cryomedia^8,9^, paraffin^8,10^, polyester wax^11^, or hard resins (Epon^12,13^, araldite^14^)—either introduce substantial artifacts that distort fine structure to the point of being uninterpretable, necessitate specimen downsizing that removes key sensory structures, or irreversibly lock tissue within the embedding matrix, precluding downstream molecular assays^11,15^. Over a century ago, nitrocellulose (celloidin) embedding^16^ became the gold standard for whole-TB histology^17–19^, preserving pristine light-microscopic morphology in intact, full-size specimens. Celloidin-based studies of the human hearing and balance organs laid the groundwork for defining their pathologies^18–21^, and later refinements enabled gentle celloidin removal for first immunohistochemical protein localization studies^22–26^—though these protocols remain applicable only in a limited manner. Overall, the “celloidin workflow” ranges from technically challenging to incompatible with multiplex immunohistochemistry^15^, RNA in situ hybridization, and sequencing-based assays^27–29^, and few, if any, studies have demonstrated these applications on celloidin-processed human TB sections. Moreover, it requires at least 13.5 months, including 2–3 weeks of fixation, up to 36 weeks of decalcification, 2 weeks of dehydration, 18 weeks of embedding, followed by serial sectioning of approximately 450 sections (20 µm thick) on heavy sledge microtomes^30^, and is both resource- and labor-intensive, with constant supply challenges for celloidin.

To overcome these barriers, we re-engineered the histology workflow for whole human TB specimens around three core objectives: (1) enhance preservation and accessibility of proteins, DNA, and RNA for molecular assays; (2) maintain celloidin-level (gold-standard) histomorphological fidelity for meaningful interpretation at tissue and cell level; and (3) reduce processing time, complexity, and cost to facilitate broad adoption, as well as streamlined section archiving and distribution. Central to our novel approach is a reversible polymethyl methacrylate (rPMMA) resin embedding workflow that leverages controlled low-temperature conditions (-40 °C to +4 °C) and eliminates months-long decalcification. After embedding, calcified specimens are sectioned near-serially (10–50 µm) via femtosecond laser microtomy or precision diamond-wire sawing, then fully deacrylated (rPMMA resin removed) from mounted tissue sections. This approach preserves exceptional histomorphology of whole human TB specimens, restores antigenicity for multiplexed immunofluorescence labeling, and preserves nucleic acid integrity for whole-genome sequencing and in situ RNA detection (e.g., RNAscope™). By addressing longstanding technical obstacles posed by human TB anatomy, our rPMMA-based workflow offers a new benchmark for multidimensional studies in human ear pathology.

## Materials and methods

### Ethics

This study was approved by the Institutional Review Boards of Massachusetts Eye and Ear (Boston, USA; #2021P001593), University Hospital Zurich (Zurich, Switzerland; #BASEC-2021-00820), and University Hospital Rostock (Rostock, Germany; #A-2020-0264). Written informed consent was obtained from all tissue donors or their legal representatives.

### Specimen harvesting and preparation

TB specimens were harvested using the intracranial bone plug technique^30^. Briefly, following craniectomy, a cylindrical bone plug containing the ear structures was removed using an oscillating autopsy saw (810 autopsy saw, Stryker, Kalamazoo, MI; 3.8 cm diameter trephine; see ^30^) (Figure 1A, harvesting). The study included seven TBs from adult donors (aged 56–78 years; postmortem interval 4–40 hours). Skeletal muscle (fresh frozen) from one donor was used as a sequencing control. Additionally, archival tissue sections from four donors, originally processed using the celloidin embedding protocol and stored in 80% ethanol, were included as controls for histomorphological analysis, DNA extraction, and RNA in situ hybridization. These specimens were selected based on matched postmortem intervals, with priority given to those with the shortest ethanol storage duration.

**Figure 1.**
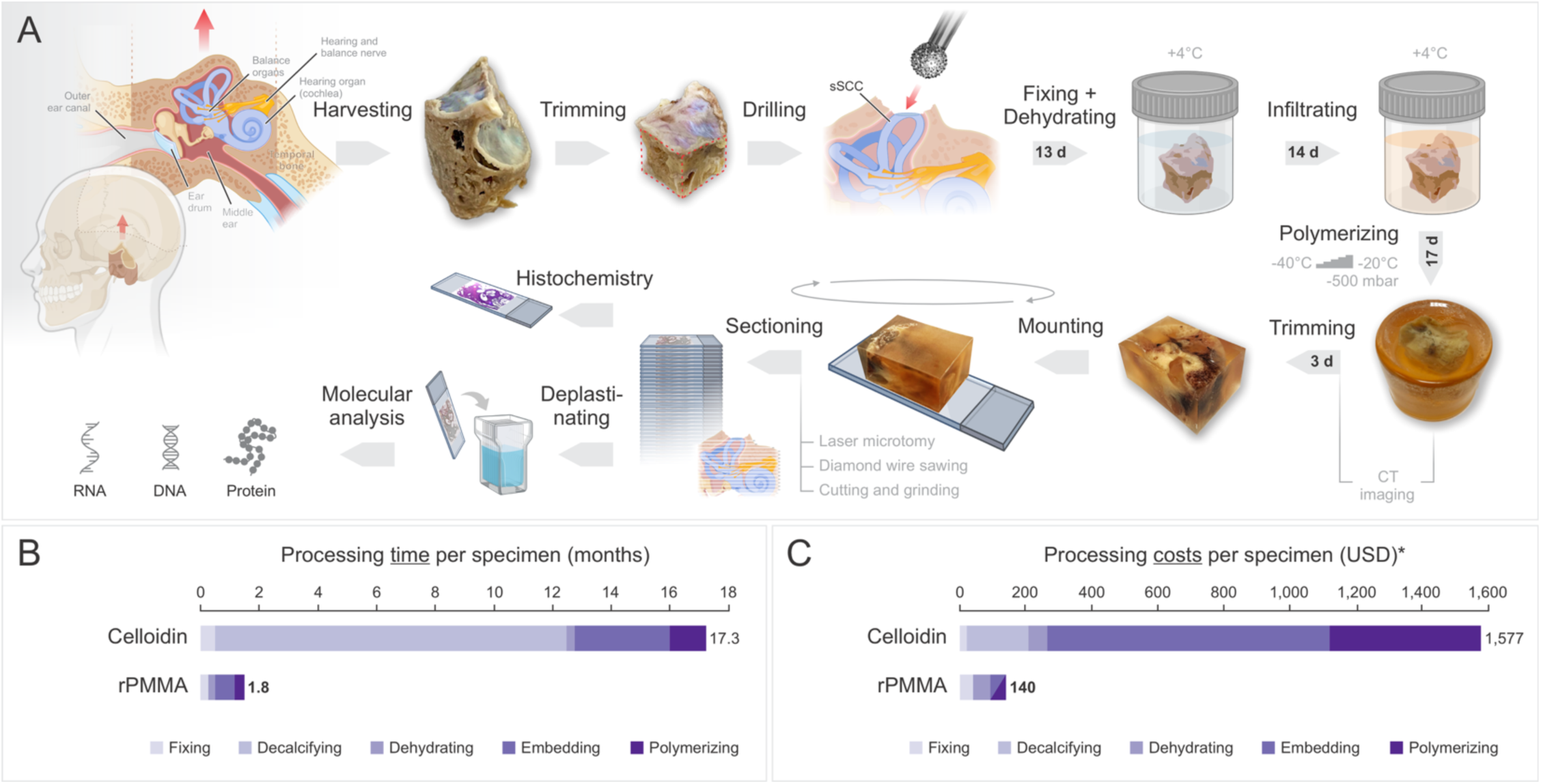
rPMMA workflow accelerates TB processing and reduces costs. (**A**) Tissue processing workflow for non-decalcified human TBs using embedding in a reversible polymethyl methacrylate (rPMMA) resin. (**B**, **C**) Comparison of processing times (B) and costs (C) between the standard celloidin protocol^30^ and the rPMMA protocol. sSCC, superior semicircular canal; *, excluding costs for sectioning.

Harvested specimens were immediately placed in ice-chilled 10% neutral buffered formalin for laboratory transfer. Excess bone on each specimen side was trimmed using a tabletop pathology bandsaw (#BBS-82203, IMEB Inc., San Marcos, CA) equipped with a diamond blade (#SD311, IMEB Inc.), reducing dimensions to approximately 35 × 25 × 25 mm (Figure 1A, trimming). Dura mater was removed, and the bone covering the superior semicircular canal was carefully drilled (#4-6 diamond burr; Figure 1A, drilling) and finally opened it with a bone curette to facilitate formalin penetration and avoid scattering bone debris from drilling into the inner ear spaces.

### Fixation and dehydration

Specimens were fixed in 10% formalin at 4°C for 6 days, then dehydrated through a graded ethanol series (30%, 70%, 80%, 95%, 100% ethanol; 24 hours each), followed by two cycles each of absolute ethanol and xylene (#X5-4, Fisher Scientific, Waltham, MA) or Histo-Clear™ (#HS-200, National Diagnostics, Atlanta, GA), 24 hours per cycle (Figure 1A, fixing + dehydrating).

### rPMMA resin embedding

Embedding of TB specimens was performed using the reversible (i.e., deacrylatable to free embedded and sectioned tissue) polymethyl methacrylate (rPMMA) resin system Technovit® 9100 (#64715444, Kulzer GmbH, Hanau, Germany, or #813-900, Ted Pella Inc., Redding, CA), following the manufacturer’s recommendations. *Preparation of solutions*: All embedding solutions were freshly prepared and stored at -20°C until use. Destabilized base solution (BS): The base solution was destabilized by passage through a chromatography column (53 mm × 457 mm; PTFE stopcock, #31-500-983, Fisher Scientific) filled with 50 g aluminum oxide (#AA4717136, Fisher Scientific). Pre-infiltration Solutions (PIS) 1: Mixture of 100 ml xylene (or Histo-Clear™) and 100 ml stabilized BS. PIS 2: 0.5 g hardener 1 dissolved in 200 ml stabilized BS. PIS 3: 0.5 g hardener 1 dissolved in 200 ml destabilized BS. Infiltration Solution (IS): 0.5 g hardener 1 and 16 g PMMA dissolved in 200 ml destabilized BS. PMMA powder was slowly added and stirred for 2–4 hours to ensure complete dissolution. Polymerization Solution A (PS-A): 0.4 g hardener 1 and 36.8 g PMMA in 230 ml destabilized BS, stirred 2–4 hours for full dissolution. Polymerization Solution B (PS-B): 2 ml hardener 2 and 1 ml regulator mixed in 25 ml destabilized BS. When not in use, all solutions were stored at – 20 °C. *Infiltration protocol* (Figure 1A, infiltrating): Specimens were sequentially incubated in PIS 1, PIS 2, and PIS 3, each for 3 days at 4°C, followed by IS for 4 days at 4°C and then at - 40°C for an additional day. During transitions between solutions, specimens were gently rotated to release trapped gas bubbles and handled with non-metal (polyoxymethylene) forceps (#BR112805, Millipore-Sigma, Burlington, MA). *Polymerization protocol* (Figure 1A, polymerizing): Polymerization was performed in a benchtop freezer modified with a 1-inch rear access port (–45 °C; #2002010, Super Coldcube, Industrial Freezer Sales, Agoura Hills, CA) using a vacuum desiccator (#F42012-0000, 0.09 cu. ft.; SP Bel-Art, Wayne, NJ) connected to a wall vacuum (–500 mbar) through this port. Polymerization solutions A and B, along with all materials were pre-cooled for at least one hour to –40°C. PS-A (90 ml) and PS-B (10 ml) were mixed in a 9:1 ratio and poured into a 100 ml glass jar (#3TRF1, Grainger, Lake Forest, IL). The specimen was carefully immersed, rotated gently to release trapped gas bubbles, and placed in the desiccator inside the benchtop freezer. Polymerization occurred at –40°C under –500 mbar vacuum for three days, with daily brief interruptions to rotate specimens and release gas bubbles. On day three, the specimen was positioned in its final orientation (tegmen down) and a 5 mm layer of paraffin oil (#MPX00461, Fisher Scientific) was added atop the polymerization solution to exclude oxygen, critical for complete polymerization. Temperature was gradually increased by increments of 5°C every three days until –15°C, maintaining a –500 mbar vacuum. After three days at –15 °C, the glass jar containing the specimen was moved to a refrigerator and held at 4 °C (without vacuum) for one further day. The paraffin oil was then drained, and the polymerized specimen was carefully freed by fracturing the jar with a hammer. See Supplementary material 1 (“Technovit® 9100 Embedding Protocol for Calcified Human Temporal Bones”) for a detailed lab version. All rPMMA-processed TB specimens were labeled sequentially as TV 1 through TV 7, according to their specimen number.

### High resolution peripheral quantitative computed tomography (HR-pQCT) imaging

TB specimens embedded in rPMMA were scanned using an XtremeCT II HR-pQCT system (Scanco Medical, Brütisellen, Switzerland). Imaging was performed at an isotropic voxel resolution of either 61 µm³ or 82 µm³. The scan parameters included an x-ray tube potential of 68 kVp, x-ray tube intensity of 147 mA, and integration time of 100 milliseconds. The acquired DICOM datasets were subsequently imported into 3D Slicer software (version 5.6.1; https://www.slicer.org). The anatomical plane representing the longitudinal axis through the cochlea’s modiolus was defined. Fiducial markers (sticky tape) indicating this mid-modiolar plane were placed directly on the surface of each embedded specimen.

### Histological sectioning

Resin blocks with embedded human TBs were first trimmed to a 25 mm maximum width to fit standard microscope slides and cut along the cochlear midmodiolar plane (see previous paragraph), using a tabletop bandsaw (#BBS-82203, IMEB Inc.) fitted with a diamond saw blade (#SD311, IMEB Inc.), and operated with water irrigation to prevent thermal damage of the resin and specimen (Figure 1A, trimming). The block’s cut surface was then smoothed using a disc sander (TG 125/E, Proxxon, Hickory, NC) with 600 grit sanding discs. Finally, each block was mounted—cut side down—onto a plain microscope slide (#12-550-10; Fisher Scientific) with a transparent, rapid-set two-component epoxy (Araldite Crystal Clear, Huntsman Advanced Materials; Figure 1A, mounting).

Sections 20–50 µm thick were prepared using one of three methods (Figure 1A, sectioning). *Laser microtomy*: Mounted blocks were positioned on the motorized stage of a TissueSurgeon device (LLS ROWIAK LaserLabSolutions, Hannover, Germany). A femtosecond laser (1030 nm wavelength, 300 fs pulse duration, 10 MHz repetition rate) was focused from beneath through a microscope slide, penetrating 20 µm into the specimen to evaporate resin and embedded tissue. Sections were produced within 10–15 minutes and subsequently polished. *Precision diamond wire sawing*: Specimen blocks were secured in a standard holder of a diamond wire saw device (DWS.100, Diamond WireTec, Weinheim, Germany). Sections were cut using a 50 µm thin diamond-coated steel wire (#DW050T, Diamond WireTec) at a maximum speed of 4 m/s, yielding approx. 50 µm thick sections within 10–15 minutes without requiring polishing. *Tabletop bandsaw and disk sander*: Blocks were guided along a straight-edge guide through tabletop band saw (#BBS-82203, IMEB Inc.) fitted with a diamond saw blade (#SD311, IMEB Inc.), producing sections approximately 300 µm thick. These were thinned and polished to a final thickness of about 50 µm using a disc sander (TG 125/E, Proxxon) with a 2000 grit sanding disc. After each cut, the blocks were remounted and the sectioning procedure repeated to generate additional 20–50 µm sections.

### Deacrylating mounted hTB sections

For subsequent immunohistochemistry, DNA extraction, or in situ RNA labeling mounted sections were deacrylated by placing the slides—tissue side down—in a glass staining jar filled with 2-methoxyethyl acetate (2-MEA; #M011325ML, Fisher Scientific) for 30 minutes, allowing the dissolved resin to drip away by gravity (Figure 1A, deplastinating). Sections were then rehydrated sequentially in xylene (or Histo-Clear™), graded ethanol solutions, distilled water, and finally PBS, each for 10 minutes.

### Archival, celloidin-processed TB sections

Celloidin-processed TB specimens, archived at Mass Eye and Ear (Boston, USA) and the University Hospital Zurich (Zurich, Switzerland) and designated sequentially as CL 1 through CL 3, were used as controls to benchmark histomorphological preservation and DNA extraction quality. Specimens were fixed in 10% neutral buffered formalin for 7–10 days, decalcified in 0.27 M ethylenediaminetetraacetic acid (EDTA; #S311-3, Fisher Scientific) for approximately 270–450 days depending on specimen size, and dehydrated in a graded ethanol series for 10 days. Embedding was performed over 84 days using increasing concentrations (up to 12%) of celloidin (nitrocellulose; #10838, LADD Research, Williston, VT) dissolved in a 1:1 mixture of absolute ethanol and ethyl ether (#E138-1, Fisher Scientific). The final embedding blocks were hardened for 14 days, followed by immersion in chloroform (#C298SK4, Fisher Scientific) and cedarwood oil (Polysciences Inc., #04851-4) for two weeks each prior to sectioning. Individual sections were mounted on numbered pieces of onion skin paper in a dish of 80% ethanol, and stored long-term in 80% ethanol at room temperature. For histomorphological comparison, archived sections from specimens most recently processed and matched for postmortem interval and donor age were selected. For DNA extraction experiments, the most recently archived sections from the University Hospital Zurich, stored for 20–21 years, were used.

### Histochemical staining

Sections from rPMMA-embedded human TBs were stained using hematoxylin and eosin (HE) or von Kossa methods without prior deacrylation, either manually or with an automatic slide stainer (Tissue-Tek Prisma, Sysmex Europe GmbH, Norderstedt, Germany; Figure 1A, Histochemistry). Stained sections were coverslipped using permanent mounting medium (Vectamount, #H-5000, Vector Laboratories, Newark, CA; or Permount, #SP-100, Fisher Scientific).

### Enzyme immunohistochemistry and multiplexed immunofluorescence labeling

Tissue sections were blocked for 15 minutes in PBS containing 5% normal horse serum (NHS) and then incubated overnight at room temperature with primary antibodies (diluted 1: — in 1% NHS; see Supplementary table 1 for antibody and fluorescent secondary combinations). After three 10-minute PBS washes, sections were incubated for 1 hour at room temperature with the appropriate secondary antibodies (also diluted in 1% NHS), followed by three additional PBS rinses. For chromogenic detection, the Fast Red Substrate Kit (#ab64254, Abcam) was applied per the manufacturer’s instructions; these sections were then dehydrated through graded ethanol, cleared in Histo-Clear, and coverslipped with Vectamount permanent mounting medium (#H-5000, Vector Laboratories, Newark, CA, USA). For multiplexed immunofluorescence, either fluorescently conjugated secondaries or biotinylated secondaries (Vectastain ABC-HRP Kit, #PK-4000, Vector Laboratories) plus a streptavidin-linked fluorescent probe were used; fluorescent sections were counterstained with Hoechst 33258 (1:1000 in PBS; #H3569, Thermo Fisher) and coverslipped in Vectashield antifade medium (#H-1000-10, Vector Laboratories). Negative controls omitted the primary antibody.

### DNA extraction, quantification, and quality assessment

DNA was extracted from individual 20 µm tissue sections (from one TB specimen per donor; samples TV 1–7). To remove the adhesive, slides were warmed to 70°C, and sections were gently detached. Each section was deacrylated in 50 mL of 2-MEA for 20 minutes under constant agitation, then rinsed three times in PBS for 5 minutes each. For celloidin-embedded sections, tissues were immersed in methanol saturated with sodium methoxide at a 1:2 dilution in methanol for 5 minutes, followed by a 100% methanol rinse at room temperature; this process was repeated three times^31^. For samples TV 1–6 and CL 1–3, DNA was extracted using the PrepFiler™ BTA Forensic DNA Extraction Kit (#4463352, Thermo Fisher Scientific, Switzerland), a kit specifically optimized for calcified tissues and those containing adhesives. DNA from sample TV 7 and its corresponding fresh frozen specimen was isolated using the QIAamp® DNA FFPE Kit (#56404, Qiagen, Germantown, MD) according to manufacturer’s instructions. DNA quantity and quality were evaluated using the human-specific Quantifiler™ Trio DNA Quantification Kit (#4482910, Thermo Fisher Scientific) alongside PicoGreen dsDNA assay kit (#P7589, Thermo Fisher Scientific) and Nanodrop spectrophotometry, ensuring consistency with established standards.

### Short tandem repeat analysis

A total of 600 pg of the extracted DNA was processed using the AmpFLSTR NGM SElect PCR Amplification Kit (#4457889, Thermo Fisher Scientific), amplifying 16 autosomal short tandem repeat (STR) loci as well as the amelogenin gene (AMEL) used for gender identification (sex-typing).

### DNA library preparation and whole-genome sequencing

For samples TV 1–6 and CL 1–3, genomic DNA was first sheared using a Covaris E220 focused ultrasonicator (#500239, Covaris, Woburn, MA). Libraries were then prepared with the NEBNext® Ultra™ II DNA Library Prep Kit (#E7645, New England Biolabs, Ipswich, MA). DNA overhangs were end-repaired, adenylated, and ligated to the NEB hairpin adapter. An adapted bead cleanup (1.8 ×) for low-input protocols was used, and the resulting libraries were amplified using unique P5 and P7 PCR primers (NEB, Ipswich, MA). Library quality and quantity were assessed with the Qubit 1.0 Fluorometer (Fisher Scientific) and Agilent TapeStation (Agilent, Waldbronn, Germany), then normalized to 10 nM in 10 mM Tris-Cl (pH 8.5) with 0.1% Tween 20. For samples TV7 and TV7-FF, DNA (1 µg input) was processed independently using the TruSeq PCR-Free library prep kit (#20015962, Illumina, San Diego, CA), following the manufacturer’s guidelines. Libraries were quantified with the Qubit dsDNA HS Assay Kit (#Q32851, Invitrogen, Waltham, MA) and sized using the BioAnalyzer 2100 High Sensitivity DNA Kit (#5067-4626, Agilent, Santa Clara, CA). They were then normalized to 4 nM based on these readings and re-quantified using a qPCR assay with the KAPA Library Quantification Kit for Illumina Libraries (#KK4824, Roche, Basel, Switzerland).

### Sequencing QC and analysis

The pooled libraries of the samples TV 1–6 and CL 1–3 were sequenced using a paired- end read configuration with 150 bp per read using the Illumina Novaseq 6000 (Illumina, Inc, San Diego, CA) and demultiplexed with bcl2fastq allowing for 1 mismatch in barcode 2. Reads were quality-checked with fastqc^32^ and were aligned to the human hg38 reference genome using bowtie2 and default parameters. For samples TV 7 and TV 7 FF, libraries were pooled and sequenced using the Illumina NovaSeq 6000 sequencing platform on a 2×150 bp paired end SP flowcell (Illumina, San Diego, CA). Sequencing data was processed and analyzed using the DRAGEN germline analysis pipeline, available through the Basespace Sequencing Hub (https://basespace.illumina.com/). DRAGEN provides end-to-end secondary analysis including mapping, alignment and variant calling (PMID: 33214604, PMID: 38260545). Sequencing reads were aligned to the human hg38 reference genome and analysis was performed using DRAGEN default parameters.

### In situ total RNA labeling and RNase A-mediated validation

Sections were labeled in situ with either Syto RNASelect (0.5 µM; #S32703, Invitrogen) or StrandBrite Green RNA Dye (10 µM; #17610, AAT Bioquest) diluted in DEPC-treated water, incubated for 30 minutes at room temperature in the dark. After three 10-minute washes in RNase-free water, nuclei were counterstained with Hoechst 33258 (1 µg/mL; #H3569, Thermo Fisher) for 15 minutes, followed by three additional washes. Confocal image stacks of the spiral ganglion region were acquired on a Leica SP5 with a 40 × water-dipping objective (NA 1.1), using sequential channels to avoid bleed-through (Syto RNASelect: 490/530 nm; StrandBrite: 488/520 nm; Hoechst: 405/460 nm). Without removing the slide from the stage, sections were then covered with either RNase A (100 µg/mL; #EN0531, Fisher Scientific) or RNase-free water and incubated for 3 hours at room temperature. The same field was re-imaged under identical settings, and loss of RNA-selective fluorescence versus stable Hoechst signal confirmed dye specificity.

### RNAscope™ in situ hybridization

Chromogenic in situ hybridization was performed using the RNAscope™ 2.5 HD Duplex Detection Kit (#322430; Advanced Cell Diagnostics, Newark, CA, USA) according to the manufacturer’s instructions. Pretreatment of 5 µm tissue sections comprised the following steps: incubation in hydrogen peroxide at room temperature for 10 minutes; heat-induced epitope retrieval in Target Retrieval Buffer in a pressure cooker within a microwave for 15 minutes; and protease Plus treatment at 40 °C for 30 minutes. Sections were then hybridized in the HybEZ Oven with the kit’s positive-control probes—peptidylprolyl isomerase B (PPIB; channel 1) and RNA polymerase II subunit A (POLR2A; channel 2). After sequential amplification, PPIB was visualized with the green chromogen and POLR2A with Fast Red, yielding discrete green and red signals, respectively. Slides were briefly rinsed, dehydrated through graded ethanols, cleared in xylene, and coverslipped with Permount mounting medium (Fisher Scientific). For detection of calbindin-2 (CALB-2) transcripts, separate serial sections underwent identical pretreatment, hybridization, and amplification using the human CALB-2 probe (channel 2; #422179 Hs-CALB2, Advanced Cell Diagnostics). The CALB2 signal was developed with Fast Red.

Following RNAscope™, sections were immunofluorescently labeled for neurofilament heavy chain (NF-h), CellMask Deep Red (#C10046, Fisher Scientific), and counterstained with Hoechst 33258. Antibody and dye dilutions, fluorophore conjugations, and incubation conditions are provided in Supplementary table 1. Finally, slides were coverslipped with ProLong™ Gold Antifade (#P36930, Fisher Scientific) mounting medium.

### Image acquisition and postprocessing

Sections stained with colorimetric (enzymatic) detection methods were imaged using a Nikon Eclipse E800 microscope (Nikon Instruments, Melville, NY) equipped with Nomarski optics and a DS-Ri2 digital color camera (Nikon Instruments) or an Olympus BX51 microscope (Olympus, Tokyo, Japan) equipped with a DP70 digital color camera (Olympus). Immunofluorescence labeled sections were imaged using a Leica TCS SP8 confocal microscope (Leica Microsystems GmbH, Wetzlar, Germany).

### RNAscope™ signal quantification

Images of primary vestibular (Scarpa’s ganglion neurons were acquired showing chromogenic CALB2 (red) and fluorescent NF-h, CellMask™ Deep Red, and Hoechst 33258 signals. In Dragonfly software (v.2024.1, Comet Technologies, Montreal, Canada), the CALB2 and NF-h channels were loaded and each converted to a binary mask using Otsu thresholding (with a lower threshold for CALB2 and a higher one for NF-h). NF-h fluorescence signal not associated with vestibular ganglion cell soma was manually removed, and individual NF-h– positive ganglion cell ROIs were defined. A Boolean “AND” operation between the CALB2 and NF-h masks then quantified the CALB2-positive pixels within each NF-h–positive soma. Finally, for each cell, CALB2-positive area was expressed as a percentage of its total NF-h–positive area.

### Statistics

Quantitative data are reported as mean and standard deviation (SD) unless otherwise specified. Normality of distribution was assessed using the Shapiro–Wilk test. For comparisons between two groups, an unpaired two-tailed Student’s t-test was used; for paired measurements of fluorescence intensity before and after RNase A treatment a paired t-test was applied. Statistical analyses were performed in GraphPad Prism 9.0 (GraphPad Software, San Diego, CA), with a two-tailed p < 0.05 considered significant. Exact n (biological and technical replicates) and p-values are indicated in the figure legends.

## Results

### Improved processing efficiency and cost savings

The novel rPMMA method (Figure 1A) dramatically accelerates and reduces the cost of human TB processing compared with the traditional celloidin workflow. Whereas celloidin processing—from initial fixation to complete embedding—required on average 17.3 months, rPMMA shortens this timeline to just 1.8 months by eliminating the prolonged EDTA decalcification step and utilizing a faster resin polymerization process (Figure 1B). Correspondingly, per-specimen costs fall from approximately USD 1,577 with celloidin to USD 140 using rPMMA (Figure 1C). These advantages are partially offset during the sectioning stage. Unlike decalcified, celloidin-embedded TBs, calcified rPMMA blocks require hard cutting techniques (see Materials and Methods), which are more labor-intensive and produce fewer sections per unit time. Nonetheless, rPMMA sections, in contrast to celloidin, are compatible with high-spatial resolution, multimodal molecular assays—including applications not possible with the traditional workflow (see subsequent paragraphs)—and thus offer the potential for substantially greater data yield and previously unattainable data dimensions (RNA level) per section, an advantage that will likely outweigh reduced sectioning throughput.

### Gold-standard equivalent histomorphological preservation of hearing and balance organs in human temporal bone specimens

To evaluate whether the rPMMA method preserves histomorphology on par with the gold-standard celloidin method, we qualitatively compared HE-stained rPMMA sections (Figure 2) to celloidin controls (Supplementary figure S1) from human TBs matched for short postmortem interval (4–7 h). At low magnification rPMMA section overview, the global tissue architecture was remarkably well preserved across the entire specimen with all anatomical regions appeared intact, undistorted, and in their native spatial relationships (Figures 2A, S1A). Closer views of the cochlear and vestibular organs demonstrated the sensory organs suspended in natural positions in their native fluid-filled compartments, including the cochlear duct (Figures 2B, S1B) and saccule and utricle in the vestibular portion (Figures 2C, S1C), matching celloidin. High-magnification images confirmed excellent cellular preservation, with clearly delineated cytoarchitecture in the organ of Corti (Figures 2D, S1D), spiral ganglion neurons (SGNs) and their axonal projections in Rosenthal’s canal (Figures 2E, S1E), and the saccular macula (Figures 2F, S1F). Sensory and supporting cells, as well as neuronal and stromal components, were clearly distinguishable, with preservation quality matching that observed in celloidin-processed specimens. Notably, unlike decalcified celloidin sections, the rPMMA method retains the inorganic (calcium carbonate) matrix of otoconia, permitting direct visualization of these biomineralized crystals within their native tissue context (Figures 2F, S1F). Together, these findings demonstrate that the rPMMA workflow achieves soft tissue preservation equivalent to the histological (celloidin) gold standard.

**Figure 2.**
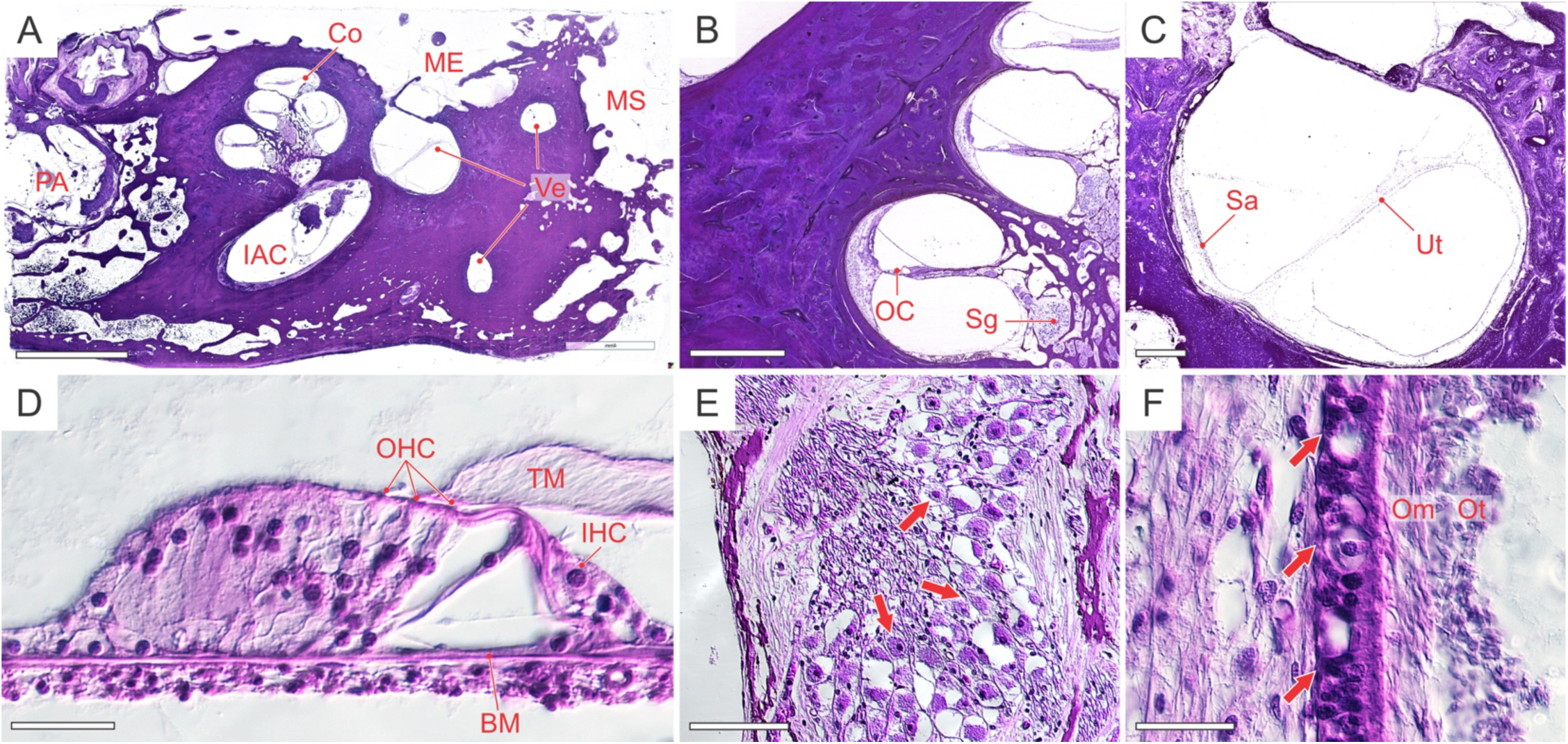
Histomorphology of hearing and balance organs in rPMMA-processed human TBs. (**A**) Overview of hematoxylin/eosin (HE) stained tissue section (Co, cochlea; IAC, internal auditory canal with cochlea-vestibular nerve; ME, middle ear; MS, mastoid space; PA, petrous apex; Ve, vestibule). (**B, C**) Details of the cochlea (B) and the vestibule (C) from (A) (OC, organ of Corti (sensory epithelium); Sg, spiral ganglion (first-order auditory neurons); Sa, saccule, Ut, utricle). (**D – F**) High-power views of Oc (D; BM, basilar membrane; OHC, outer hair cells; IHC; inner hair cells; TM, tectorial membrane), Sg (E; arrows, neuron cell bodies), and Sa (F; arrow, macular epithelium; Om, otoconial membrane; Ot, calcified otoconia (compare to Supplementary figure S1F where Ot are missing). Scale bars: 5 mm (A), 500 μm (B and C), 50 μm (D, E, and F).

### Preservation of native cochlear implant–tissue interface

To test whether rPMMA embedding combined with hard cutting techniques—i.e. femtosecond laser microtomy and precision diamond wire sawing—maintains the native cochlear implant-tissue interface, we processed a human TB with a cochlear implant electrode array in situ (Figure 3A). Examination of the polymerized block’s cut surface showed the silicone-sheathed electrode array in the basal and middle cochlear turns, supported by the rigid resin matrix (Figure 3B). Contact-free femtosecond laser sectioning and abrasive diamond wire sawing then produced sections that preserved the electrode array’s genuine positioning within the scala tympani (Figure 3C). At high magnification, multinucleated giant cells—key mediators of the inner ear’s foreign-body response^33,34^—were seen lining the silicone sheath, demonstrating preservation of intricate interface details (Figure 3D). Previous histology workflows leveraged by our laboratory for human TBs carrying metal implants could not achieve this level of preservation. In the celloidin method, the soft nitrocellulose scaffold cannot support direct cutting of the electrode with steel or tungsten-carbide blades, forcing electrode removal before sectioning and obliterating the tissue–implant interface (Figures S2A–B). Araldite embedding permits cutting with tungsten-carbide blades but still produces displacement and shattering artifacts at the interface (Figures S2C–D) and is incompatible with many downstream molecular assays. To our knowledge, this is the first demonstration of hard-resin embedding paired with advanced sectioning methods that preserves undisturbed cochlear implant-tissue contacts in human temporal bone while remaining compatible with molecular analyses.

**Figure 3.**
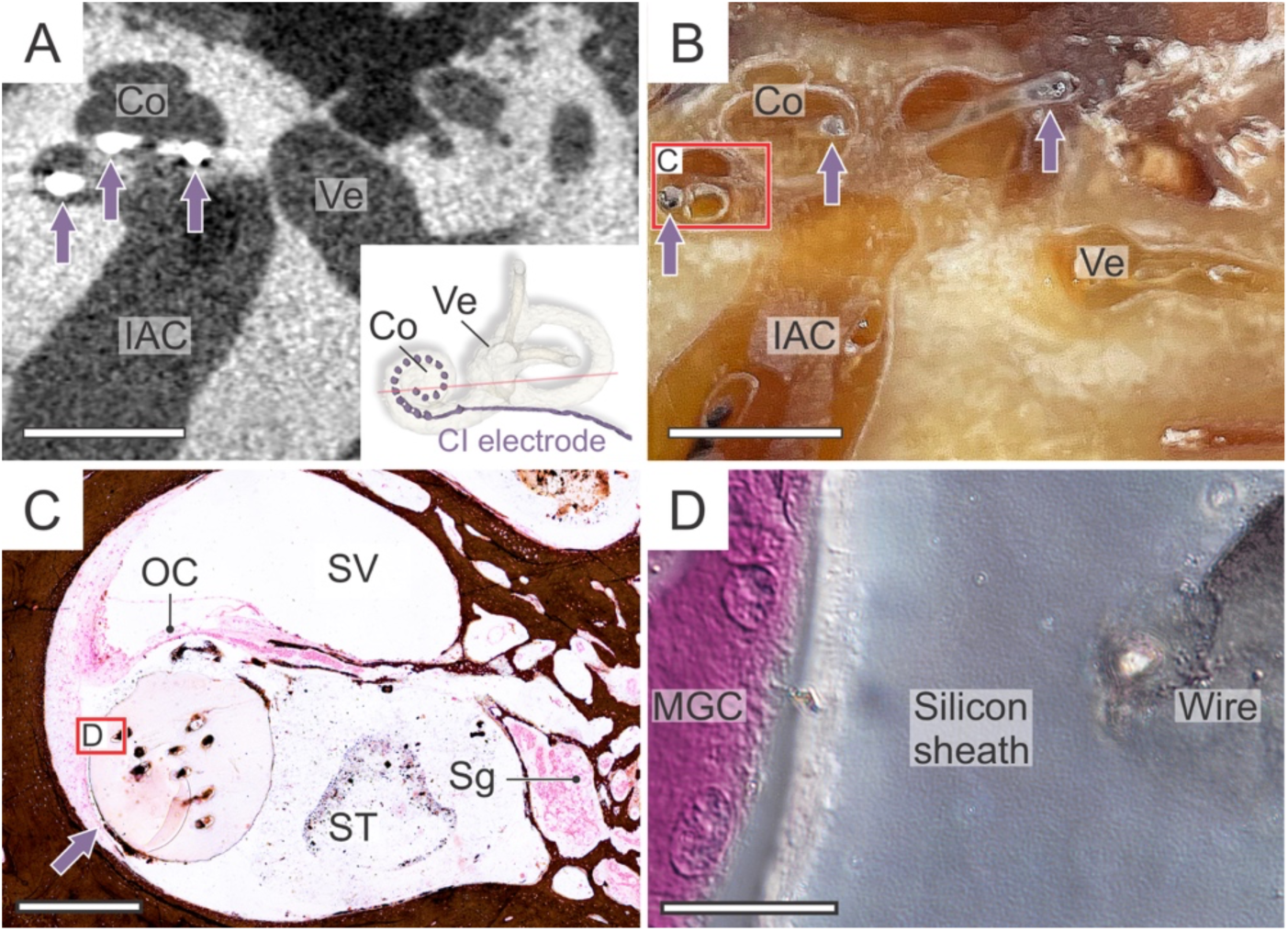
rPMMA-embedded human TB with cochlear implant electrode in situ. (**A**) Axial CT of rPMMA block with embedded TB showing cochlear implant electrode (arrows) in basal and middle turns of the cochlea (Co). Inset: 3D reconstruction of Co, cochlear implant electrode, and vestibular labyrinth (Ve); red line indicates CT imaging plane. IAC, internal auditory canal. (**B**) Cut-surface view of rPMMA block with the embedded TB from (A). Arrows indicate cochlear implant electrode. (**C**) Cross-section of the basal cochlear turn with CI electrode (arrow) in scala tympani (ST). Von Kossa stain highlights calcium-rich bone matrix in brown. SV, scala vestibuli; OC, organ of Corti; Sg, spiral ganglion. (**D**) High-power view of implant-tissue interface (location indicated in (C)) showing a multinucleated giant cell (MGC) abutting the electrode’s silicon sheath. Scale bars: 5 mm (A, B), 500 μm (C), 50 μm (D).

### Improved compatibility with immunohistochemistry and multiplex immunofluorescence

To assess the compatibility of rPMMA-processed human TB sections with antibody-based labeling, we performed chromogenic and multiplexed immunofluorescence labeling following deacrylation. Enzyme-based (chromogenic) immunohistochemistry for osteopontin (OPN) yielded strong, specific labeling of bone matrix within the mineralized otic capsule (Figure 4A), confirming antigen accessibility in the calcified and deacrylated tissue section. In adjacent sections from the same specimen, NF-h robustly labeled SGN cell bodies (Figure 4B). Multiplex immunofluorescence of a cochlear duct cross-section with antibodies against alpha-smooth muscle actin (α-SMA) and NF-h (Figure 4C) specifically labeled hair cell stereocilia and auditory nerve fibers in the organ of Corti (Figure 4C, inset), while CellMask and Hoechst 33258 highlighted the exceptionally well-preserved cytoarchitecture in the deacrylated sections. In triple-antibody labeling experiments, we observed strong, specific fluorescence: OPN intensely labeled the extracellular bone matrix, dentin matrix protein 1 (DMP1) specifically marked osteocytes, and type I collagen (Coll I) delineated the fibrous tissue matrix adjacent to the calcified otic capsule (Figure 4D). All negative controls (primary antibodies omitted) were confirmed negative (data not shown). Together, these data demonstrate that deacrylated rPMMA sections of calcified human temporal bone retain epitope integrity across both mineralized and soft tissues. Although we did not conduct a head-to-head comparison with celloidin-processed specimens, our experience shows that immunohistochemistry protocols optimized for FFPE tissues translate reliably to rPMMA-processed sections—unlike with celloidin-processed specimens.

**Figure 4.**
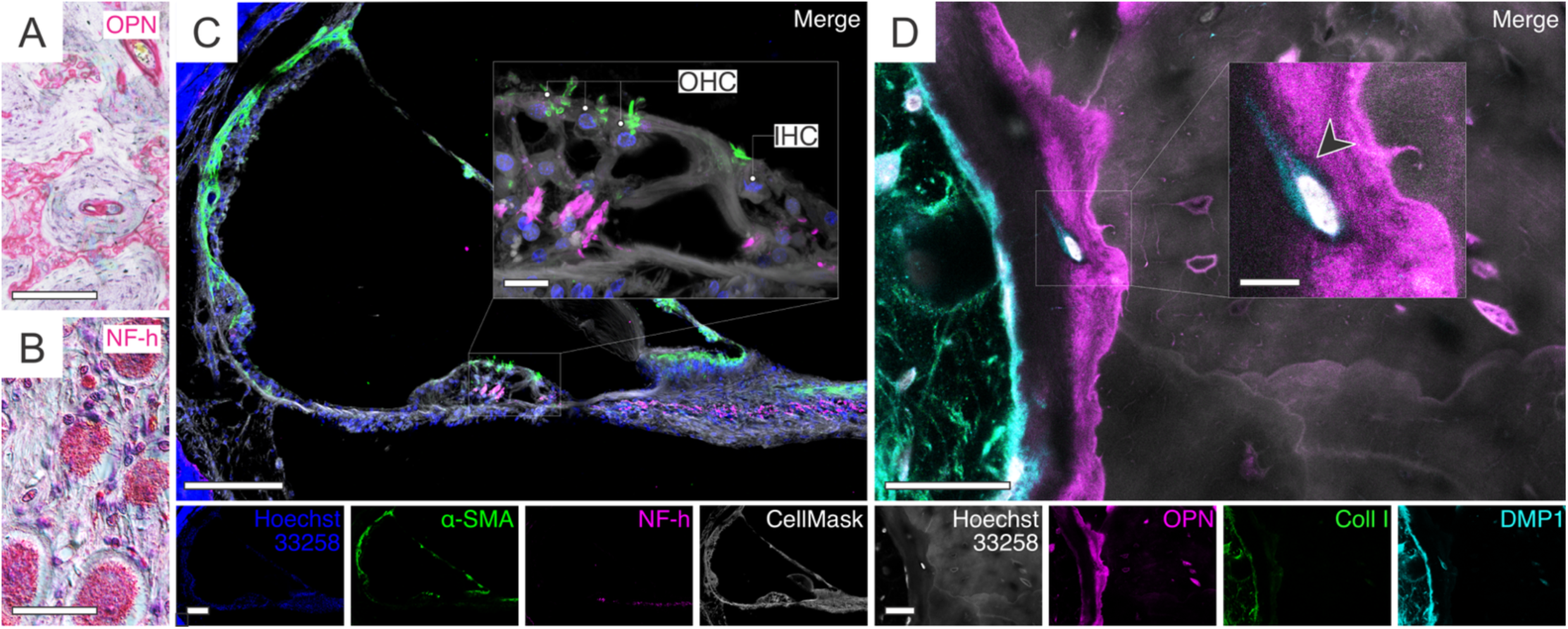
Immunohistochemistry in rPMMA-embedded human TB sections. (**A, B**) Enzyme (alkaline phosphatase)-immunohistochemistry for osteopontin (OPN) outlines structures (globuli interossei) in otic capsule bone (A); neurofilament-heavy (NF-h) labels spiral ganglion neuron cell bodies (B); nuclei counterstained with HE. (**C**) Immunofluorescence for alpha-smooth muscle actin (α-SMA; green), NF-h (magenta), CellMask (white), and Hoechst 33258 (blue) in a cochlear duct cross-section; inset shows organ of Corti with OHC and IHC. (**D**) Immunofluorescence for OPN (magenta), collagen I (Coll I; green), and dentin matrix protein-1 (DMP1; cyan) with Hoechst 33258 (blue) in the calcified otic capsule bone and adjacent fibrous tissue. Scale bars: 200 µm (A), 50 µm (B), 200 µm (C), 50 µm (D), 10 µm (insets, C and D).

### High-yield recovery of intact genomic DNA and compatibility with whole-genome sequencing

To assess DNA recovery from rPMMA-processed tissue sections, we extracted genomic DNA from single sections of six human TB specimens with postmortem intervals ranging from 4 to 40 hours (mean 16.8 h, SD 6.7 h). The average yield was 15.76 ng per section (SD 18.13 ng; Figure 5A), with a mean degradation index of 5.6 (SD 2.5; Figure 5B) and DNA integrity number (DIN) of 2.85 (SD 0.25; not shown). A section with a 40-hour postmortem interval yielded only 1.63 ng of highly degraded DNA. Three celloidin-processed archival control sections yielded negligible DNA amounts (mean 0.005 ng, SD 0.004 ng) and were excluded from downstream analysis. Short tandem repeat (STR) profiling showed full amplification of all 17 loci in 60% of rPMMA samples and 16 of 17 loci in the remaining 40%, with no allelic dropout or cross-human contamination (Figure 5C), demonstrating that the rPMMA matrix both preserves native DNA integrity and protects against exogenous (e.g., laboratory staff) DNA contamination by fully sealing the tissue within the polymerized block. Minimal bacterial contamination was detected (mean 0.47% of reads; SD 0.40%; Figure 5D). These results demonstrate that rPMMA embedding preserves sufficient DNA integrity and yield for downstream molecular analyses in specimens with postmortem intervals below 24 hours.

**Figure 5.**
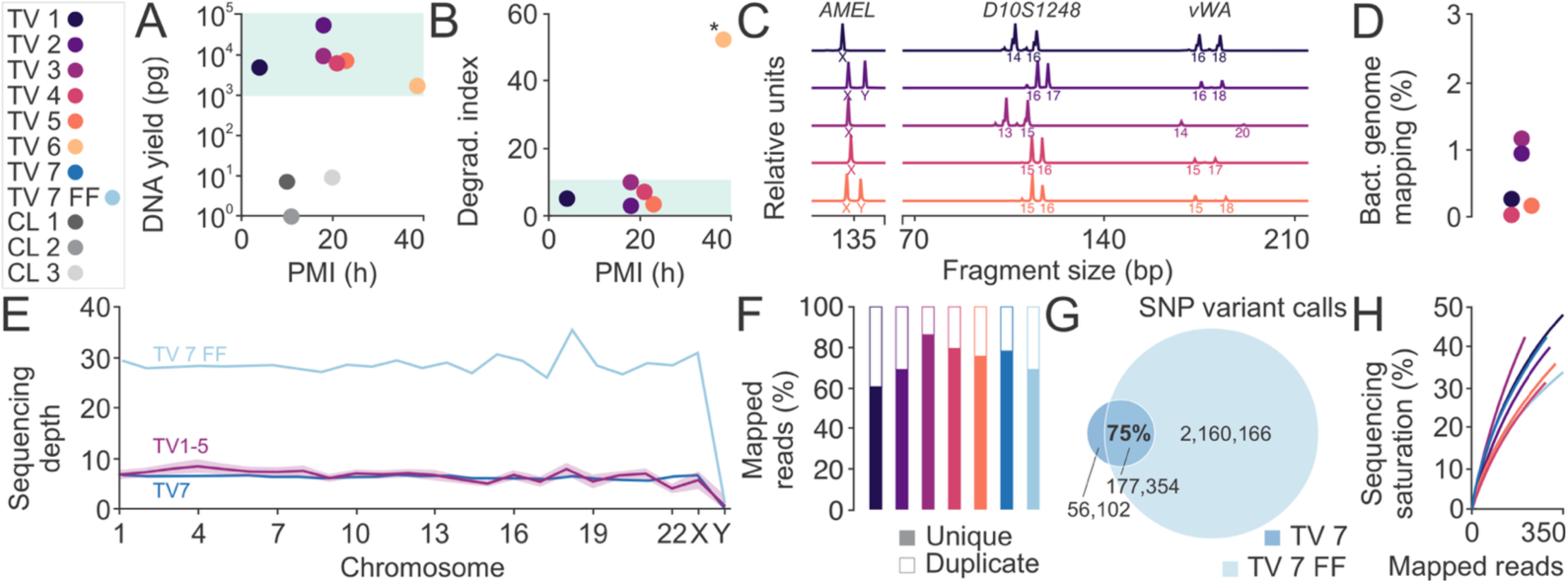
DNA Extraction and WGS metrics from rPMMA-vs. celloidin-embedded human TBs. (**A–B**) DNA yield (A) and integrity (B) in relation to the postmortem interval (PMI) *, TV 6 (PMI 40 h) was excluded due to extensive DNA degradation. (**C**) Representative short tandem repeat (STR) profiles at at the X-chromosomal AMEL locus and autosomal D10S1248 and vWA loci. (**D**) Bacterial genome mapping for TV 1–5. (**E**) Average sequencing coverage across the genome: TV 1–5 (purple), TV 7 (blue), and a fresh-frozen (FF) control from the same donor as TV 7 (cyan). (**F**) Sequencing saturation curves based on unique mapped reads. (**G**) Counts of unique versus duplicate mapped reads for each sample. (**H**) Sequencing saturation curves for TV 1–7 and TV 7 FF.

To determine whether DNA extracted from single rPMMA-embedded sections is suitable for whole-genome sequencing (WGS), we performed WGS directly on DNA from individual tissue sections. Each sample generated approximately 400 million reads, achieving a mean coverage depth of 7.37 × (SD 0.70; Figure 5E), with an average duplication rate of 30.8% (SD 8.3%; Figure 5F), consistent with expectations for low-input DNA. Comparison of single nucleotide polymorphism (SNP) variant calls from a matched rPMMA section (TV 7) and a fresh frozen sample (TV 7 FF) from the same specimen showed 75% concordance (Figure 5G), indicating that most true variants are retained despite some dropouts. Saturation analysis indicated a continued increase in variant detection without reaching a plateau for all samples, suggesting that deeper sequencing would further improve yield (Figure 5H). These data confirm that single rPMMA-processed human TB sections are fully compatible with WGS, delivering reliable variant detection and scalable genomic coverage.

### Preservation of total RNA species

To evaluate the preservation and in situ accessibility of total RNA species in rPMMA-processed human TB sections—and to compare this with the best-case archival celloidin sections—we performed fluorescent labeling with two RNA-selective dyes (Syto RNASelect and StrandBrite Green). Specimens were matched by postmortem interval (9–10 hours), formalin fixation (6–7 days), and archival storage of less than six months, with celloidin sections kept in 80% ethanol and rPMMA sections stored dry and dark at room temperature. In rPMMA sections, Syto RNASelect yielded nuclear, nucleolar, and cytoplasmic signals in SGNs, a pattern distinctly different from the co-labeled DNA-specific dye Hoechst 33258 (Figure 6A). In contrast, SGNs in celloidin sections exhibited only Hoechst-positive nuclei, with no detectable Syto RNASelect fluorescence (inset, Figure 6A), indicating marked loss of tissue RNA species. To confirm the specificity of the RNA signal in rPMMA tissue, we carried out an on-slide RNase A digest: sections were co-labeled with StrandBrite Green and Hoechst and imaged to establish baseline fluorescence (Figure 6B–B′), then incubated on the microscope stage for 3 hours at room temperature in either RNase A or water (photobleaching control), and finally re-imaged (Figure 6C–C′). After correcting for photobleaching, RNase A treatment caused a significant decrease in nuclear StrandBrite fluorescence (Figure 6B″–B‴), while Hoechst intensity remained statistically unchanged (Figure 6C″–C‴), thereby confirming the in situ specificity of RNA-selective staining in rPMMA-embedded human TB sections.

**Figure 6.**
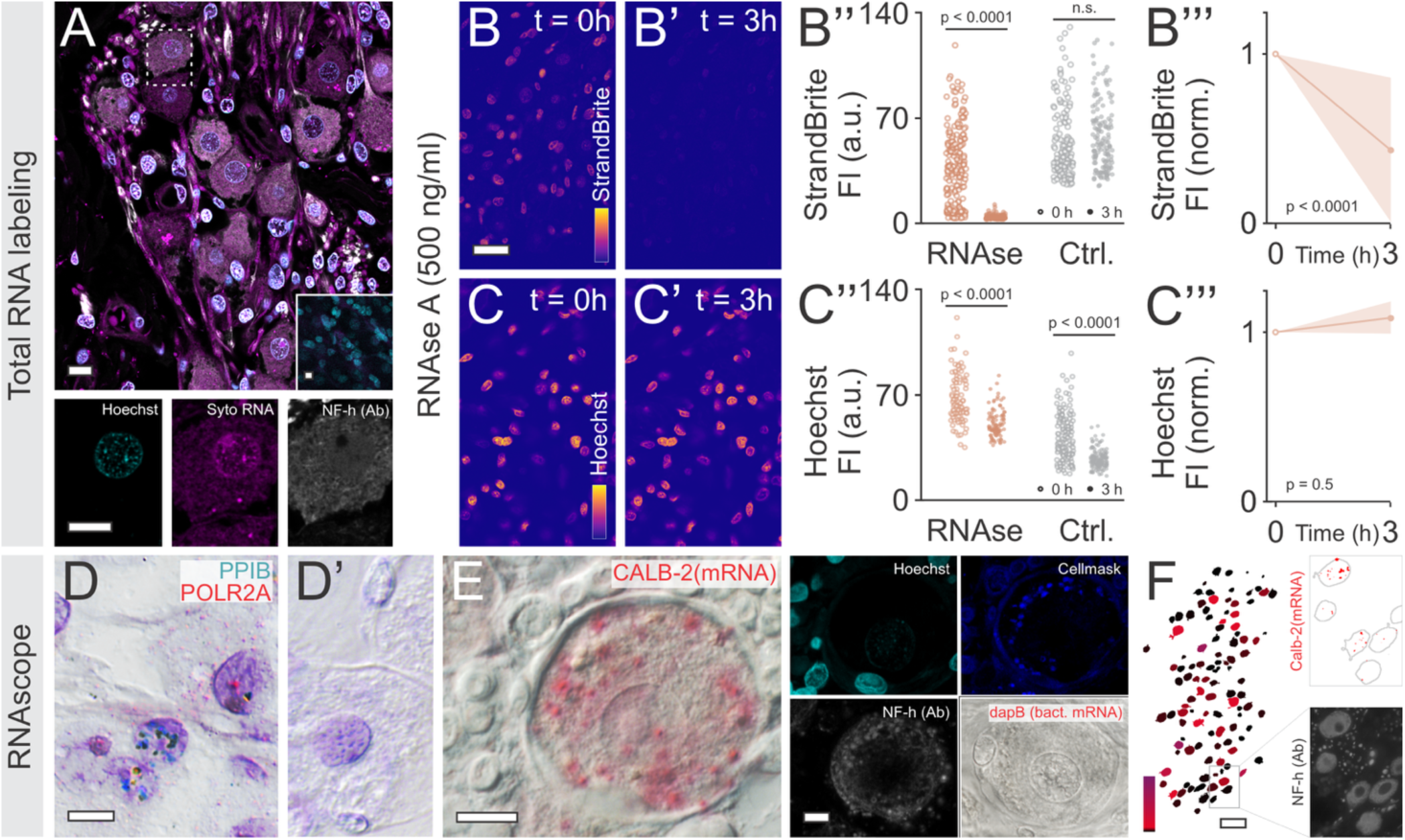
Total RNA in situ labeling and RNAscope™ in rPMMA-versus celloidin-processed human TBs. (**A**) Syto RNAselect (RNA-selective; magenta) and Hoechst 33258 (DNA-selective; blue) fluorescent labeling in rPMMA-embedded spiral ganglion cells (SGNs), with NF-h (white). Inset: SGNs in celloidin section. No Syto RNAselect labeling observed. (**B-C’’’**) StrandBrite (RNA-selective; B–B’’’) and Hoechst 33258 (C–C’’’) in SGNs at 0 min (B, C) and after 3 h (B’, C’) digestion with RNase A (500 ng/ml) versus control (DEPC-water only). Fluorescence intensity (FI) for StrandBrite (B’’) and Hoechst 33258 (C’’), and corresponding decay slopes (B’’’, C’’’). (**D, D’**) Chromogenic RNAscope™ targeting housekeeping mRNAs PPIB and POLR2A in SGNs (D) and bacterial (dabP) negative control (D’). (**E, F**) Chromogenic RNAscope™ for Calbindin-2 (CALB-2) mRNA (E, red) combined with fluorescent NF-h (green), CellMask™ membrane stain, and Hoechst (blue); bacterial negative control inset. (F) Quantification of Calb-2–positive area in 100 NF-h⁺ vestibular ganglion neurons. Scale bars: 10 µm (A), 20 µm (B–C’), 10 µm (D–E), 100 µm (F).

### Targeted mRNA integrity confirmed via RNAscope™

We assessed whether the rPMMA workflow and subsequent deacrylation preserve mRNA in human TB sections sufficiently for high-resolution in situ hybridization. Using RNAscope™, we first applied positive controls—PPIB and POLR2A—and a bacterial DapB probe as a negative control. In rPMMA sections, PPIB yielded punctate nuclear signal in non-neuronal cells, while POLR2A signal localized to both nuclei and cytoplasm of SGNs (Figure 6D). The bacterial DapB probe produced no detectable signal (Figure 6D′), confirming assay specificity. We then probed for human CALB2 mRNA. Fast Red chromogenic puncta were observed within SGN cell bodies (data not shown) and prominently in vestibular ganglion neurons co-labeled for NF-h, CellMask, and Hoechst (Figure 6E). Quantitative image analysis—segmenting each vestibular ganglion neuron by its NF-h–defined boundary and calculating the fraction of area occupied by CALB2 puncta—revealed marked heterogeneity in CALB2 transcript abundance across individual neurons (Figure 6F). Together, these data demonstrate that rPMMA-processed human TB sections retain mRNA integrity and in situ accessibility, enabling reliable, single-cell–level detection of target transcripts by RNAscope™.

## Discussion

Here, we introduce an innovative histology workflow for calcified human TBs that combines embedding in a rigid PMMA resin with near-serial thin-sectioning via femtosecond laser microtomy and precision diamond-wire sawing. This approach preserves gold-standard morphological detail—on par with the classic celloidin method—while remaining fully compatible with both in situ and sequencing-based molecular assays. By compressing a process that traditionally exceeds 17 months into about six weeks, and by eliminating toxic reagents and high costs, this method provides an unprecedented platform for fully integrated, spatially resolved histo-molecular studies of human auditory and vestibular tissues.

Hearing and balance disorders pose a substantial, growing global health burden: over 430 million people live with disabling hearing loss, a figure set to exceed 700 million by 2050^35^, chronic middle ear infections affect hundreds of millions of children each year^36^, and dizziness and vertigo afflict up to 20 percent of adults annually, with a lifetime prevalence around 7–8%^37^. Yet in-depth investigation of hearing and balance organ pathologies in the human ear is severely constrained by existing methods—a critical barrier to advancing disease understanding and effective therapies^4^. The century-old, widely adopted celloidin histology workflow preserves exquisite morphology but is largely incompatible with in situ protein and nucleic acid assays^31^; standard formalin-fixation paraffin-embedding (FFPE) supports many molecular assays^38,39^, but markedly distorts the human ear’s microanatomy, rendering cellular-level interpretation unreliable^24^; and dissociative molecular approaches (e.g. single-cell RNA sequencing) generate rich cell-specific expression profiles, yet sacrifice the three-dimensional tissue architecture critical for spatial contextual insight.

Our rPMMA workflow addresses these challenges through several key innovations. Its methacrylate chemistry crosslinks primarily with itself and interacts minimally with endogenous proteins and nucleic acids, preserving biomolecules for downstream analysis. Once cured, the hard, refractive-index–matched resin preserves global and cellular architecture, and immobilizes metallic implants in place. Precision sectioning via contactless femtosecond laser microtomy or abrasive diamond-wire sawing then yields clean, artifact-free slices through both tissue and metal without displacement or distortion. The reversible PMMA matrix can be readily removed with solvent, exposing the tissue for robust, multiplexed immunofluorescence labeling; high-yield, high-quality DNA extraction—while effectively preventing foreign DNA contamination, a common issue with celloidin^28^ and paraffin^40^. Library-complexity metrics— including non-saturated sequencing saturation curves and high overlap of unique fragments compared to fresh-frozen controls—demonstrate that even at the modest exploratory read depths we chose, our WGS libraries remain rich in novel content, highlighting the potential for sensitive calling of rare single-nucleotide variants, structural rearrangements, and mosaic mutations. We also successfully performed total RNA in situ labeling and targeted RNAscope for abundant mRNAs, providing proof-of-principle for gene-specific in situ assays on rPMMA-processed human TB sections. Although factors such as postmortem interval, RNA degradation kinetics in different TB tissues, and residual resin removal conditions will need systematic study to define which specimens are amendable for RNA studies, this workflow broadly paves the way for high-resolution, large-scale spatial transcriptomics of human auditory and vestibular tissues. The entire protocol completes in under two months and reduces per-specimen costs by over 90%, enabling truly integrated, spatially resolved histo-molecular investigations of postmortem human auditory and vestibular tissues. Rather than serially sectioning entire specimens, our workflow focuses on producing a select few high-quality sections that each yield rich, high-information molecular data.

During method development, we identified three key limitations: (i) *Outgassing during polymerization*. PMMA’s exothermic free-radical curing can release dissolved gases and residual monomer, generating microscopic bubbles that distort surrounding tissue. We reduced this by using a very slow, multi-stage thermal ramp (from –40 °C to –10 °C, finishing at 4 °C), which mitigates gas evolution during solidification. However, bubbles still form within the bony labyrinth cavities, where they cannot escape. (ii) *Section adhesion and solvent resistance*. Deacrylation requires aggressive solvents (here 2-MEA) that quickly dissolve standard histology glues and dislodge sections. We solved this by mounting 20 µm slices with a two-component Araldite epoxy: it cures to a crystal-clear, solvent-resistant bond that endures hours of solvent exposure. However, its long (hour-scale) cure time limits throughput. Faster UV-curable adhesives may accelerate slide preparation in the future. (iii) *Sectioning throughput and accessibility*. Earlier approaches applying diamond sawing to calcified human TBs used wide-blade band saws to cut fixed specimens into millimeter-thick slabs, thereby increasing surface area to accelerate the subsequent decalcification process^41–43^. By contrast, we use 50–80 µm diamond-coated wires to produce thin, ready-to-use sections, and completely bypassing the decalcification step. However, diamond-wire sectioning—and femtosecond laser microtomy— both require specialized equipment and considerable hands-on time, resulting in low-to-moderate throughput. Consequently, these methods are best applied to targeted regions of interest—are less suited to produce (near)serial sections of entire human temporal bone specimens—and deliver single sections that are amendable for high–information–yield molecular assays.

## Conclusion

Here we describe a histology workflow for postmortem human temporal bones that combines gold-standard morphological preservation with versatile, on-section protein and nucleic acid assays—all within a streamlined, cost-effective protocol. This approach establishes a new standard for human tissue spatially resolved molecular studies of hearing and balance disorders using human tissue.

## Acknowledgements

We thank Barbara J. Burgess and Diana T. Jones for their decades of technical expertise and stewardship of the human temporal bone collection at Mass Eye and Ear. A.H.E. is grateful to Marcus Müller (University of Tübingen) for early inspiration and invaluable discussions on Technovit® 9100. We thank Daniela Meir and Fabian Baron for their support in harvesting temporal bones during autopsies. This work was funded by the National Institute on Deafness and Other Communication Disorders (NIDCD) National Human Ear Resource Network (U24-DC0208499. Additional support was provided by the National Institute of Arthritis and Musculoskeletal and Skin Diseases (NIAMS; P30 AR075042) and an investigator-initiated research grant from MED-EL (Innsbruck, Austria), which had no involvement in study design, data collection, analysis, interpretation, or manuscript preparation.

## Supplementary figures and figure legends

**Supplementary figure S1.**
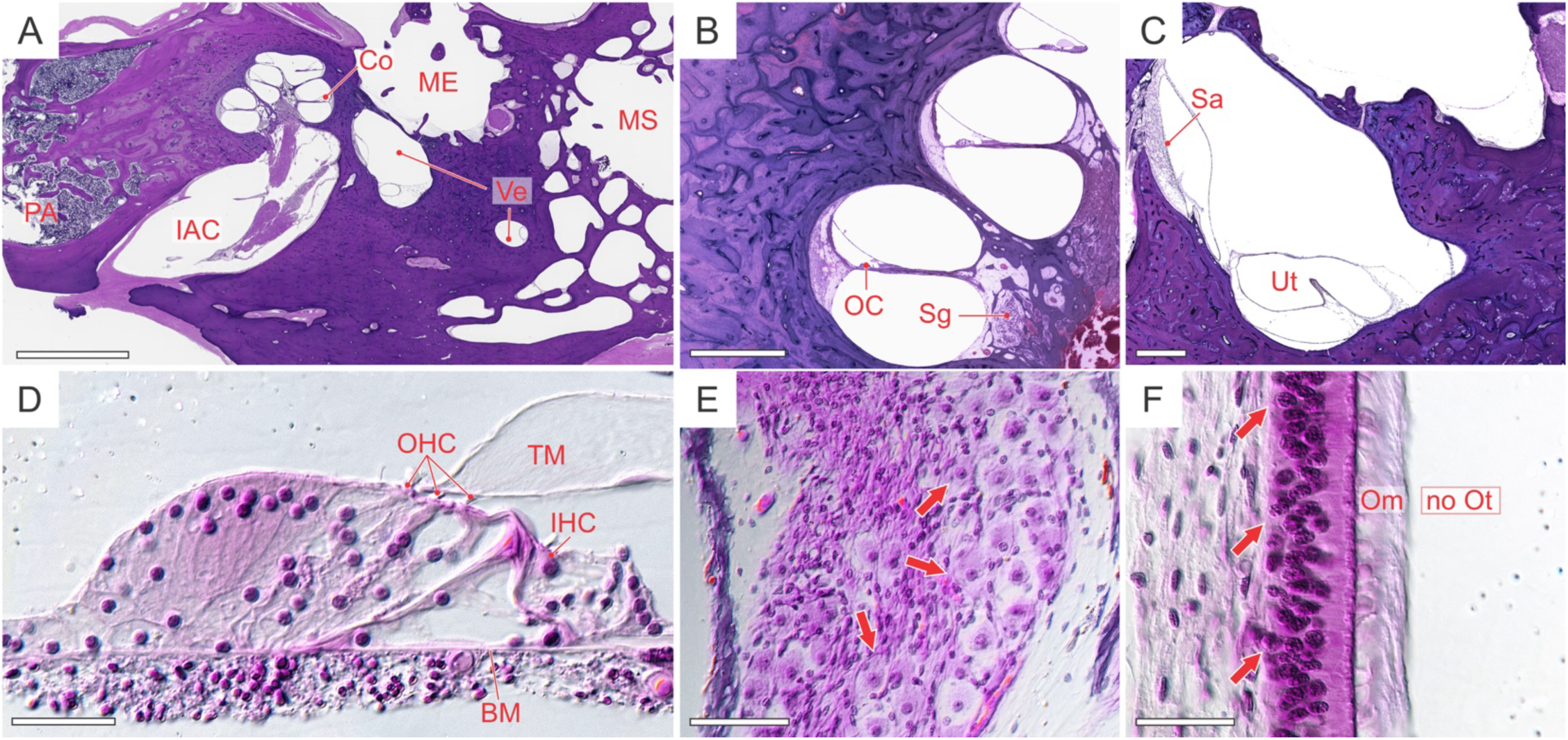
Histomorphology of hearing and balance organs in celloidin-processed human TBs. (**A**) Overview of hematoxylin/eosin (HE) stained tissue section (Co, cochlea; IAC, internal auditory canal with cochlea-vestibular nerve; ME, middle ear; MS, mastoid space; PA, petrous apex; Ve, vestibule). (**B, C**) Details of the cochlea (B) and the vestibule (C) from (A) (OC, organ of Corti (sensory epithelium); Sg, spiral ganglion (first-order auditory neurons); Sa, saccule, Ut, utricle). (**D – F**) High-power views of Oc (D; BM, basilar membrane; OHC, outer hair cells; IHC; inner hair cells; TM, tectorial membrane), Sg (E; arrows, neuron cell bodies), and Sa (F; arrow, macular epithelium; Om, otoconial membrane; no Ot, otoconia missing due to prolonged decalcification (compare to Figure 1F). Scale bars: 5 mm (A), 500 μm (B and C), 50 μm (D, E, and F).

**Figure S2.**
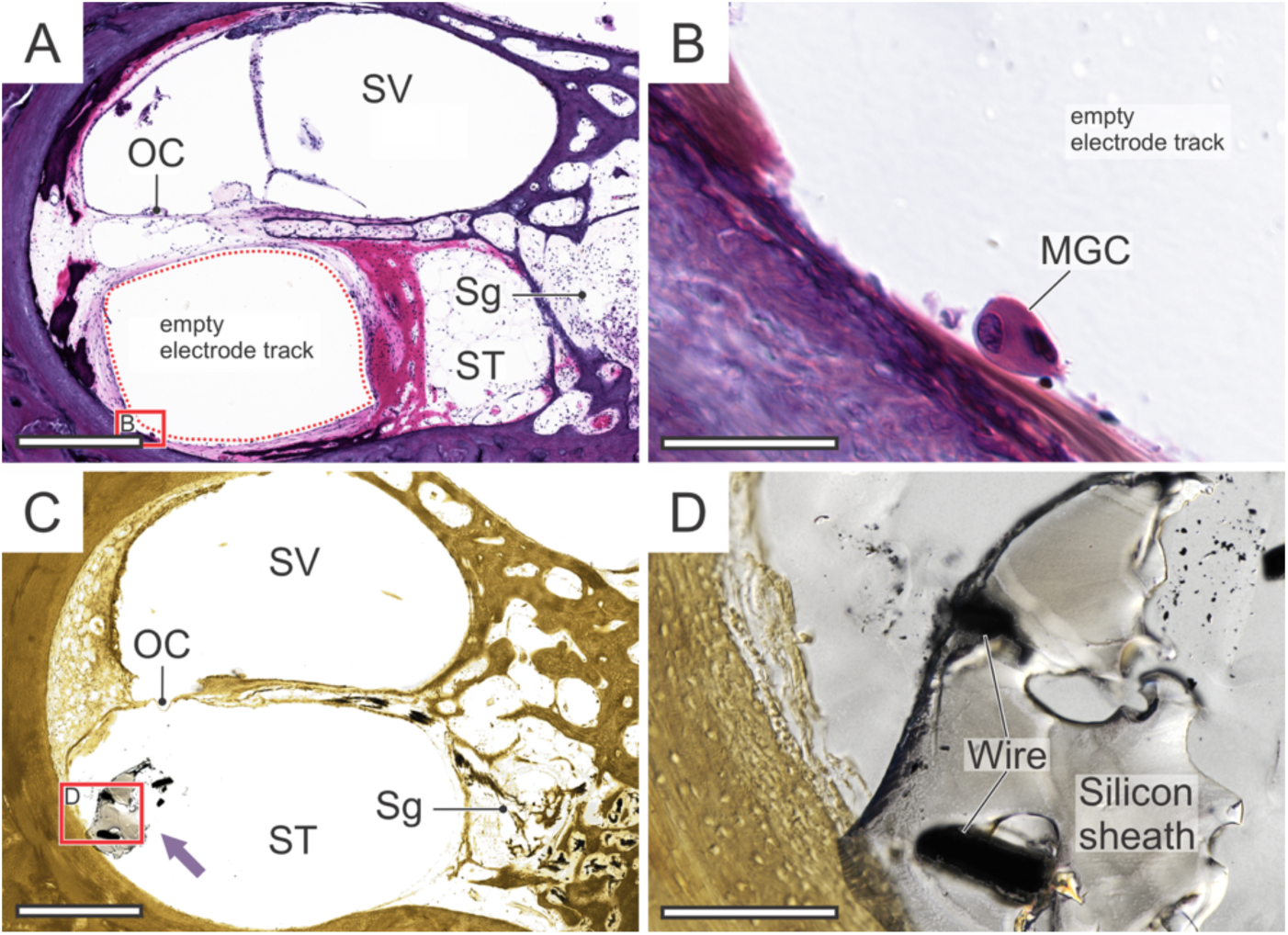
Human TBs processed in celloidin (A, B) or araldite (C, D). In celloidin-embedded specimens the CI electrode was removed prior to decalcification, yielding an empty track in the scala tympani (ST) of the basal cochlear turn: (**A**) low-power cross-section showing ST and scala vestibuli (SV) (OC, organ of Corti; Sg, spiral ganglion); (**B**) high-power view of the empty track with a multinucleated giant cell (MGC). In Araldite-embedded specimens the electrode remained in situ: (**C**) cross-section of the basal turn with the CI electrode (arrow) in ST; (**D**) high-power view showing electrode damage and displacement caused by steel-blade microtome sectioning. Scale bars: 50 µm (A, C); 50 µm (B, D).

**Supplementary table S1.** Primary and secondary antibodies used for immunostaining.

| Primary AB | Working dilution | Cat. #, Vendor | Linked secondary AB (+ tertiary link) | Working dilution | Cat. #, vendor | Figure |
| --- | --- | --- | --- | --- | --- | --- |
| Rabbit anti-osteopontin | 1 :50 | SC21742, Santa Cruz | KA-50F, Universal Alkaline Phosphatase Immunostaining Kit (for mouse and rabbit primary antibodies), Diagnostic Biosystems |  |  | 4A |
| Rabbit anti-neurofilament heavy | 1 :160 | N4142, Sigma |  |  |  | 4B |
| Mouse anti-alpha smooth muscle actin | 1 :200 | A2547, Millipore Sigma | Donkey anti-mouse IgG (H+L) Alexa 488 | 1 :400 | 715-545-150, Jackson ImmunoResearch | 4C |
| Rabbit anti-neurofilament heavy | 1 :500 | N4142, Millipore Sigma | Donkey anti-rabbit IgG (H+L) Alexa 568 | 1 :400 | 711-575-152, Jackson ImmunoResearch | 4C |
| Rabbit anti-osteopontin | 1 :250 | ab8448, Abcam | Donkey anti-rabbit IgG (H+L) Alexa 568 | 1 :400 | 711-575-152, Jackson ImmunoResearch | 4D |
| mouse anti-collagen I |  | ab21286, Abcam | Donkey anti-mouse IgG (H+L) Alexa 488 | 1 :400 | 715-545-150, Jackson ImmunoResearch | 4D |
| Sheep anti-dentin matrix protein 1 | 1 :100 | AF4386, R&D Systems | Biotinylated donkey anti-sheep IgG (H+L)<br>+<br>Streptavidin Alexa 647 | 1 :400<br>1 :800 | 713-065-147, Jackson ImmunoResearch<br>016-600-084, Jackson ImmunoResearch | 4D |

## Supplementary material 1: Technovit® 9100 Embedding Protocol for Calcified Human Temporal Bones

1. *Personal protective equipment (PPE) & safety procedures*

- Lab coat, nitrile gloves, and safety goggles
- Handle volatile or caustic chemicals (formalin, Histo-Clear™, all Technovit® 9100 Kit solutions) in a certified fume hood.
- Use face shield and cryogenic gloves when handling solutions or specimens at temperatures below –20 °C.
- Have appropriate spill kits readily available.
- Follow institutional safety and chemical waste disposal protocols; adapt as needed.
2. *Reagents & Solutions*

- Fixatives & Solvents

◦ 10% neutral buffered formalin or 4% paraformaldehyde
◦ Ethanol (50%, 70%, 80%, 95%, 100% [200 proof])
◦ Histo-Clear™ (orange oil–based xylene substitute)
- Embedding Kit

◦ Technovit® 9100 Embedding Kit (Ted Pella, Inc. #813-900)
◦ Heavy paraffin oil (Supelco PX0046-1)
- Chromatography & Destabilization

◦ Aluminum oxide (Al₂O₃), 99% (Fisher Scientific AA4717136)
◦ Chromatography column with PTFE stopcock (Fisher 31-500-983)
3. *Preparing Technovit Solutions*

- Destabilized Base Solution (dBS):

◦ Fill chromatography column with 50 g Al₂O₃ powder.
◦ Add ∼710 mL base solution (BS) to column.
◦ Collect ∼700 mL filtrate (= dBS) at ∼1 drop/s into a brown glass bottle.
- Pre-infiltration Solutions:

◦ PIS 1: 100 mL BS + 100 mL Histo-Clear™
◦ PIS 2: 200 mL BS + 0.5 g Hardener 1
◦ PIS 3: 200 mL dBS + 0.5 g Hardener 1
- Infiltration Solution (IS):

◦ 200 mL dBS + 0.5 g Hardener 1 + 16 g poly methyl methycrylate (PMMA) powder
- Polymerization Solutions (PS):

◦ PS-A: 230 mL dBS + 0.4 g Hardener 1 + 36.8 g PMMA
◦ PS-B: 25 mL dBS + 2 mL Hardener 2 + 1 mL Regulator
- Storage: Keep all solutions at –20 °C until use. Equilibrate at +4 °C before use.
4. *Specimen Processing Schedule*

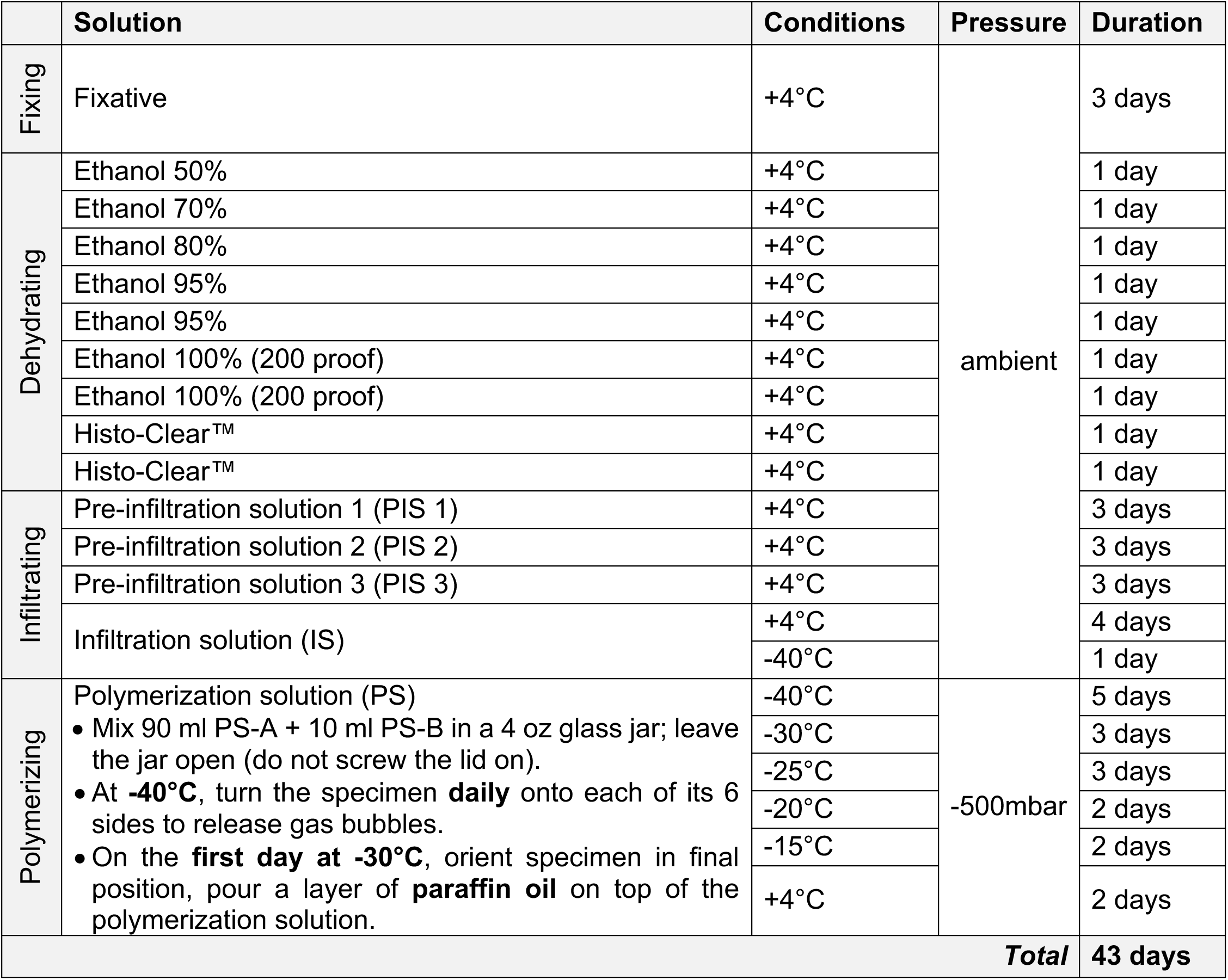
5. *Detailed Polymerization Steps*

- Chill specimen in IS at –40 °C (final day of infiltration).
- Pre-cool PS-A, PS-B, glassware, and vacuum desiccator to –40 °C.
- In fume hood, mix 90 mL PS-A + 10 mL PS-B in a pre-chilled jar; place open in the desiccator.
- Transfer specimen into mixed PS; turn on all six faces daily at –40 °C to release bubbles; re-evacuate to ∼–500 mbar.
- From day 6 onward (–30 °C stage), leave sealed; do not open until polymerization completes.
- At end of –15 °C stage, test surface hardness; if still soft, cap jar and store at +4 °C for 2 days.
6. *Post-Embedding Workflow*

- Wrap glass jar with fully polymerized resin in cloth or large piece of paper, using safety glasses safety gloves and in a container, carefully fracture glass with e.g. a hammer and extract embedded block.
- Polymerized blocks can be stored at room temperature.
- Continue with block trimming and then thin sectioning using precision diamond wire sawing or laser microtomy.

